# Biostimulation methods based on chemical communication improve semen quality in male breeder rabbits

**DOI:** 10.1101/2024.04.30.591951

**Authors:** Paula R. Villamayor, Uxía Yañez, Julián Gullón, Pablo Sánchez-Quinteiro, Ana I. Peña, Juan J. Becerra, Pedro G. Herradón, Paulino Martínez, Luis A. Quintela

## Abstract

Biostimulation aims to optimize reproductive parameters as part of animal management practices by modulating animal sensory systems. Chemical signals, mostly known as pheromones, have a great potential in this regard. This study was conducted to determine the influence of short-term male rabbit exposure to different biological secretions, potentially pheromone-mediated, on reproductive parameters of males. Four groups of 18 males each were exposed to A) doe urine, B) 2-phenoxyethanol, C) doe vaginal swab, and D) distilled water (control), three times over a 2.5h exposure window, just before semen collection. Semen volume, sperm concentration and motility, as well as subpopulation analysis of the spermatozoa were assessed for each condition. Additionally, testosterone levels in blood samples were monitored at five time points over the 2.5 h exposure window. We found a higher percentage of motile, progressive, fast progressive and mid-progressive spermatozoa in any of the three experimental groups compared to the control group. In contrast, the semen volume and the percentage of immotile and non-progressive spermatozoa was generally higher in the control group. We then identified a higher proportion of a subpopulation of fast and progressive spermatozoa in groups A, B, and C compared to group D. Our data indicates that sperm motility increases when animals are exposed to specific biological fluids potentially containing pheromones, and that an increase in sperm volume does not correlate with an increase in spermatozoa concentration, progressiveness, and speed. Finally, no differences in testosterone levels were found among comparisons, although males of groups A and C (exposed to natural female biological fluids) showed a tendency towards higher testosterone levels. In conclusion, our results indicate that rabbit sperm quality increases upon exposure to the biological secretions proposed, thereby supporting further investigation into their molecular identity. This exploration could eventually pave the way for implementing the use of pheromones in rabbit husbandry to enhance reproductive and productive parameters in farmed rabbits.

## 1. Introduction

The utilization of artificial insemination (AI) in rabbit farms for commercial purposes is relatively recent compared to its longstanding use in other species such as cattle or swine. The widespread adoption of AI in rabbit farming began in the late 80s, and it has been continuously evolving. Indeed, its introduction allowed the development of band management, with the subsequent intensification of the cycle and further enabling genetic selection for increased prolificacy [1–3]. AI is performed either with fresh or cooled semen, generally provided by semen production centres in pools from different males [4,5]. Therefore, the use of AI maximizes the profits of rabbit farming by reducing the number of pen-raised male breeders and, consequently, the number of non-productive cages [6,7]. However, the reliable evaluation of the semen and the fertilizing ability of the bucks are extremely important to achieve successful AI [8].

There are several factors that can influence qualitative and quantitative sperm production, such as collection frequency, lighting programmes, age and health status, and feeding strategies. According to the literature, a collection frequency of two ejaculates per week (obtained at least 15 minutes apart) results in good semen production [9,10]. Moreover, a daily constant programme of 16 h of light and 8 h of darkness increases sperm production compared to a shorter light duration [10,11]. In addition, semen quality (concerning concentration, progressiveness, and speed of spermatozoa) and health status decreases in older (> 2 years) rabbits [10]. Finally, there are certain feeding strategies that should be considered to ensure adequate sperm production. Males should be preferentially fed *ad libitum*, with diets that contain more than 15% of crude protein [10,12]. Similarly, fat content should be controlled, and it has been reported that balanced fatty acid compositions seem to be more important than the total amount [13]. Moreover, the high lipid unsaturation of the spermatozoa membrane makes them very susceptible to peroxidation, which endangers both the membrane’s structure and DNA integrity. Thus, the antioxidant protection is provided by the seminal plasma, and this is strongly affected by dietary supplementation [10].

Despite these methods have been shown to increase reproductive efficiency, they have not considered how animals would perform in a more natural environment. Although this perspective is challenging to achieve in intensive farming systems, the existence of biostimulation methods opens new possibilities for further increasing productivity while enabling individuals to exhibit more natural behaviours, thereby ensuring animal welfare while implementing a more organic approach.

Biostimulation constitutes a natural animal husbandry practice targeting the optimization of reproductive parameters by modulating animal sensory systems. Specifically, chemical communication, often referred to as pheromone-mediated communication, has been postulated to exert a pivotal role in biostimulation owing to its impact on sexual behaviour and reproductive physiology [14]. Most mammals use pheromones to convey information among themselves [15], and these intraspecific cues are essential for individuals and species survival. Pheromones are released into the environment by individuals through biological secretions (e.g., urine, vaginal fluid, etc.) [16–18] or exocrine glands (e.g., chin gland, lacrimal gland, etc.) [19–21], triggering a specific reaction in another individual of the same species. Due to the species-specific condition and the broad molecular nature of pheromones, studies on their chemical nature have been scarce in the literature [22]. Indeed, although certain chemicals have been implicated in biostimulation effects, most of them have not yet been conclusively identified as pheromones. An example in rabbits is 2-phenoxyethanol, identified as a fixative chemical compound of chin gland secretions of male rabbits, that is synthesized in higher concentration as the male’s dominance level increases. This substance is thought to reduce the rate at which other volatile components are released, thus prolonging the presence of a dominant animal’s scent in the environment [23]. Additionally, 2-phenoxyethanol has been identified as pheromone in insects [24].

Despite the potentiality that pheromones hold for being utilized as powerful biostimulators in male rabbit farming, most scientific studies related to biostimulation in rabbits to date have used behavioural approaches based on male-female interactions. For example, some authors have speculated that stimulating male rabbits by exposing them to females just before semen collection (SC) could increase sperm concentration [25]. A later study confirmed that exposure of males to females increases libido and also the quality and quantity of ejaculate [26].

However, to the best of our knowledge, no studies have directly exposed male rabbits to female biological compounds as potential sources of pheromones. Thus, it remains unclear which biological secretions (e.g., female urine, vaginal fluid) and which specific compounds or molecules could be responsible for such reproductive improvements. Conducting studies that directly expose animals to these secretions would facilitate the identification of specific compounds involved in chemical communication and contribute to establishing biostimulation as a routine animal practice.

Consequently, the hypothesis of our study was that female urine, vaginal fluid, and 2- phenoxyethanol, as potential sources of pheromones, could be responsible for an improvement in the reproductive performance of male rabbits. The objective of the present study was to determine if the exposure to these stimuli before SC increases the libido and semen quality and quantity of male breeder rabbits.

## 2. Materials and methods

### 2.1. Animals

This study was conducted according to the regulations and general recommendations of the National Board of Agriculture on the use of animals for scientific purposes. Seventy-two male breeder rabbits (Hyplus strain PS 40, Grimaud Frères, Roussay, France), between 45 and 163 weeks old and 5 to 7 kg in weight, were included in the study. All the procedures were carried out under farm conditions in the industrial rabbit farm COGAL SA (Rodeiro, Spain). Animals were individually housed and kept in their own cage for the entire experiment. A forced ventilation system was used, and the inside temperature was maintained between 18 °C and 22 °C using an air conditioned-heater system. Light intensity was 70 lux, with an artificial lighting program of 12 h of light and 12 h of darkness, and feeding and drinking was provided *ad libitum*. SC was performed once a week, following the farm protocols.

### 2.2. Experimental design

A behavioural experiment in the seventy-two male breeders was performed to determine whether libido, semen quantity and quality, and testosterone levels vary upon exposure to different potential sources of pheromones. We established four male groups, A, B, C and D, based on the specific compound they were exposed to: (A) Doe urine, (B) 2-phenoxyethanol (potential pheromone released by dominant males [23]), (C) Vaginal swab, and (C) Distilled water, used as a control (Table 1). Each of the four groups was composed of 18 individuals, six of which underwent blood extraction for testosterone measurements. All groups were housed in the same room, at opposite locations and facing a ventilation system, which assured no cross contamination upon odour exposure while maintaining similar environmental conditions (Figure 1A). Animals were exposed three times to the corresponding stimuli as follows: 149 min, 75 min, and 45 min before SC (Figure 1B). For groups A, B and D, the corresponding stimulant was sprayed around the nose. Specifically, 1 mL nasal spray was used in each exposure per animal (a total of 3 mL per individual and stimulant). For Group C, animals were exposed to the vaginal swab (direct contact between the swab and the nose of the animal) during ∼ 10 seconds –some animals licked it–. In any case, no direct contact between humans and animals was made to avoid stress and changes in reproductive parameters due to animal manipulation.

**Figure 1.**
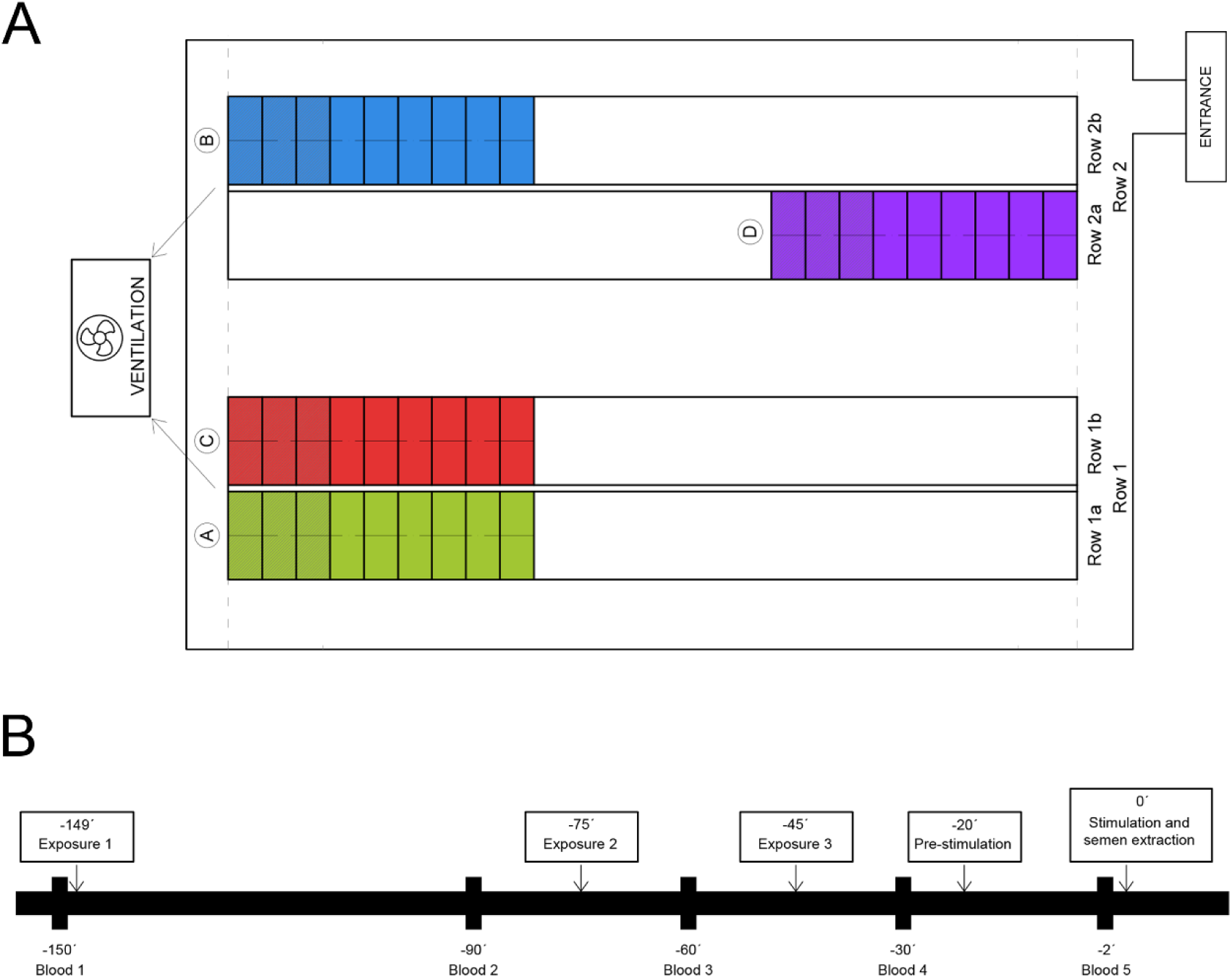
Experimental design. A) Map of the farm where the experiment was conducted. All four groups were kept in the same room to ensure similar environmental conditions. The room was organized into two rows (1,2), each with two sub-rows (1a,1b, and 2a,2b, respectively). In each sub-row, there were 50 male rabbits, divided in 25 and 25, with one subrow above the other. The four groups (A, B, C, D) were distributed as indicated. Briefly, 18 animals per group, 9 above other 9 as indicated by the dashed line drawn for each group, were used. Six individuals in each group were used for blood extraction (darker squares at the edges of each group). All groups were located strategically, facing a forced ventilation system, and therefore avoiding the circulation of stimuli across the room. B) Timeline of the experiment, showing the times for blood extraction, the three time-points of exposure to the given stimulant, and the times of pre-stimulation, stimulation, and semen collection. All times are expressed with a minus (-) that decreases as the timeline progress until “time 0”, which indicates the time of semen collection.

**Table 1:** Summary of the four different experimental groups, the type of exposure and its role in reproduction.

| Group name | Exposure | Application | Function |
| --- | --- | --- | --- |
| A | Doe urine | Directly sprayed on the nose area | Source of semiochemicals |
| B | 2-phenoxyethanol | Directly sprayed on the nose area | Potential female semiochemical |
| C | Vaginal swab | Contact between the swab and the nose of the animal | Source of semiochemicals |
| D | Distilled water | Directly sprayed on the nose area | Control |

Additionally, according to the farm protocol, a pre-stimulation and stimulation periods were developed before SC to increase male performance. This was achieved by exposing individuals to a male conspecific, that did not belong to any experimental group. This practice was done twice, 20 min (pre-stimulation) and just before (stimulation) SC, by adding the novel male to the individual’s cage for 10 seconds (Figure 1B).

### 2.3. Sample collection

The 2-phenoxyethanol solution was prepared by diluting 2-phenoxyethanol (PHR 1121, Sigma-Aldrich, St. Louis, USA) to a concentration of 1% in 25% ethanol. Urine and vaginal fluid were collected from mature does (Hyplus strain PS19), aged 6-18 months and weighing 3.5–5 kg. Does were housed separately from the males on a different farm.

Pools of 60 mL of urine were collected by ultrasound-guided cystocentesis, 24 h before the onset of the experiment, and kept at 4 °C overnight. Cystocentesis was the only method technically possible to ensure farm conditions and therefore cues delivered in the lower urinary tract might be missed.

Additionally, dry swabs were introduced in the vaginal region of multiparous, non-lactating and no-white vulvar colour females (females with white vulvar colour show lower receptivity levels [14]) for 15 seconds. A maximum of two swabs were taken from each female to ensure impregnation of vaginal fluid. Swabs containing blood or urine were discarded to avoid interference with any compound other than vaginal fluid.

### 2.3. Semen collection and processing

Semen was collected with a pre-warmed artificial vagina just after stimulation (Fig. 1B). The time between the introduction of the second male (for the stimulation) and mounting was always registered by the same person and considered an indicator for libido [26]. After SC, the total volume obtained from each animal was measured following gel removal, if present, and 100 µl semen / individual were rapidly diluted (1:40) in commercial diluent (mra-bit® standard, KUBUS LAB S.A., Madrid, Spain), stored at 16 °C and carried to the laboratory for its analysis within the following 8 h.

The sperm concentration was determined manually using a NeuBauer Chamber, and motility was evaluated using the CASA system (Sperm Class Analyzer 6.1.0; Microptic, Barcelona, Spain). Images were taken from 5 µL of the diluted semen, which were placed on a Makler chamber (Israel Electrooptical Industry, Rehovot, Israel). Three to eight microscopic fields (total spermatozoa count > 500) were analysed per sample using a microscope (Eclipse E200, Nikon, Tokyo, Japan) supplied with a prewarmed stage at 37 °C and a negative phase-contrast objective at a magnification of X 100. The settings used for the analysis were: minimum area = 25 pixels; maximum area = 100 pixels; immotile spermatozoa < 17 μm/s; slow spermatozoa: 17-25 μm/s; medium spermatozoa: 26-50 μm/s; rapid spermatozoa > 50 μm/s; progressive spermatozoa STR (percentage of straightness) > 70. Objects incorrectly identified as spermatozoa were minimized on the monitor by using the playback function. Additionally, the kinematic parameters recorded for each spermatozoon were: curvilinear velocity (VCL, µm/s), the average path velocity of the sperm head along its actual trajectory; straight-line velocity (VSL, μm/s): the average path velocity of the sperm head along a straight line from its first to its last position; average path velocity (VAP, μm/s): the average velocity of the sperm head along its average trajectory; percentage of linearity (LIN, %): the ratio between VSL and VCL; percentage of straightness (STR, %): the ratio between VSL and VAP; wobble coefficient (WOB, %): the ratio between VAP and VCL; mean amplitude of lateral head displacement (ALH, μm): the average value of the extreme side-to-side movement of the sperm head in each beat cycle; and beat cross frequency (BCF, Hz): the frequency with which the actual sperm trajectory crosses the average path trajectory [27].

### 2.4. Blood collection and testosterone determination

A total of six individuals in each of the four groups were used for blood sample collection (24 individuals in total), and five samples were collected per individual from the central auricular artery over a 2.5 h period (-150 min, -90 min, -60 min, -30 min, and just before (∼ 2 min) SC; ∼ 1 mL per timepoint). The first blood sample was collected from males just before their exposure to the stimulants to serve as a control for physiological hormonal levels. The rest of the blood samples were collected between exposures (Figure 1B). Blood was left at room temperature for 30 to 45 minutes and then centrifuged twice at 5500 rpm for 10 minutes. The obtained blood serum was aliquoted and immediately kept in dry ice and stored at -80 °C until further analysis. Testosterone concentration was determined using a Rabbit Testosterone ELISA Kit (Cusabio Technology LLC, Houston, TX, USA), following the manufacturer’s instructions.

### 2.5. Statistical analysis

A total of sixty seven male breeder rabbits were included in the statistical analysis – five animals were not considered due to incidences during sample collection. Descriptive statistics for the response time, semen volume, concentration, and motility parameters (percentage of immotile, motile, progressive, rapid progressive, medium progressive, and non-progressive spermatozoa) within each group were recorded. A Kruskal-Wallis test was carried out to check for differences among the experimental groups corresponding to the abovementioned variables.

Concerning T, a General Linear Model (GLM) repeated measures test was performed including the collection times as intra-subject variables and the group as covariable. Additionally, a multivariate k-means cluster analysis was carried out to classify the motile spermatozoa into four clusters according to their kinematic parameters. The k-means clustering model used euclidean distances computed from the 8 quantitative variables, after standardization of the data (arcsine transformation), so that the cluster centres were the means of the observations assigned to each cluster. The specified number of clusters was based on the previous analysis of hierarchical dendograms [28] constructed on individual rabbits using the Ward method. Contingency tables were used to determine the percentages of spermatozoa assigned to the different motile subpopulations within each group. Thereafter, a Kruskal-Wallis test was performed to determine differences regarding sperm subpopulations between groups. All analyses were conducted in SPSS version 28.0 for Windows (SPSS Inc., Chicago, IL, USA). Differences were considered significant at p ≤ 0.05.

## 3. Results

### 3.1. Libido evaluation and sperm analysis

Our results showed statistically significant differences between the experimental groups A, B, C, and group D regarding the percentage of immotile, motile, progressive, rapid progressive, and medium progressive spermatozoa (p < 0.05). These results indicate that exposure to the given biological fluids might have increased the sperm motility of rabbit male breeders compared to control individuals. Additionally, the percentage of non-progressive spermatozoa significantly differed between Group D and groups A and C (p < 0.05). Regarding volume, differences were observed between Group A and groups C and D (p < 0.05). Similarly, the sperm concentration obtained for groups A and C was significantly different from Group D (p < 0.05). No differences were observed among groups for the response time (p > 0.05). The results for the descriptive statistics and the Kruskal-Wallis test regarding the response time and the sperm analysis are displayed in Table 2.

**Table 2:** Results for the Kruskal-Wallis test (Mean ± SD) including seventy seven semen samples, obtained from breeder rabbits that were exposed to female urine (Group A), 2- phenoxyethanol (Group B), vaginal swab (Group C) or distilled water (Group D), preserved at 16°C and analysed within 8 h after collection using the CASA system.

| Parameters | Experimental group |  |  |  |
| --- | --- | --- | --- | --- |
|  | (A)Female urine | (B)2-phenoxyethanol | (C)Vaginal swab | (D)Distilled water |
| VOL (mL) | 0.94 ± 0.29<sup>a</sup> | 1.02 ± 0.26<sup>ab</sup> | 1.23 ± 0.42<sup>b</sup> | 1.38 ± 0.58<sup>b</sup> |
| Concentration (M/mL) | 1173.47 ± 400.32<sup>a</sup> | 897.78 ± 328.07<sup>ab</sup> | 1112.22 ± 332.45<sup>a</sup> | 797.50 ± 466.55<sup>b</sup> |
| Response time (S) | 2.96 ± 2.32<sup>a</sup> | 3.03 ± 2.09<sup>a</sup> | 3.89 ± 3.57<sup>a</sup> | 3.02 ± 1.05<sup>a</sup> |
| Total motility (%) | 70.15 ± 23.50<sup>a</sup> | 49.15 ± 33.93<sup>a</sup> | 51.80 ± 32.14<sup>a</sup> | 22.65 ± 23.69<sup>b</sup> |
| Total progressiveness (%) | 55.06 ± 22.16<sup>a</sup> | 39.20 ± 30.16<sup>a</sup> | 39.23 ± 27.81<sup>a</sup> | 15.97 ± 19.04<sup>b</sup> |
| Rapid progressive motility (%) | 21.86 ± 11.26<sup>a</sup> | 19.14 ± 17.76<sup>a</sup> | 17.39 ± 12.08<sup>a</sup> | 7.17 ± 9.01<sup>b</sup> |
| Medium progressive motility (%) | 33.20 ± 14.16<sup>a</sup> | 20.06 ± 13.87<sup>b</sup> | 21.84 ± 17.10<sup>b</sup> | 8.81 ± 10.51<sup>c</sup> |
| Non-progressive motility (%) | 15.09 ± 4.27<sup>a</sup> | 9.95 ± 4.69<sup>b</sup> | 12.57 ± 6.79<sup>a</sup> | 6.68 ± 5.80<sup>b</sup> |
| Immotile spermatozoa (%) | 29.85 ± 23.50<sup>a</sup> | 50.85 ± 33.93<sup>a</sup> | 48.21 ± 32.14<sup>a</sup> | 77.35 ± 23.69<sup>b</sup> |

*Different superscript letters indicate significant differences between groups ( $p < 0.05$ ). VOL: volume; S: seconds.*

### 3.2. Sperm subpopulation analysis

We then performed a sperm subpopulation analysis for a more complete evaluation. From the seventy-seven ejaculates evaluated, kinematic parameters of 36,819 motile spermatozoa were determined, and four subpopulations with different motility patterns were identified by the multivariate analysis. Means (± SD) of each kinematic parameter defining the four subpopulations are displayed in Table 3. Specifically, subpopulation 1 (SP1) included 24.27% of the total motile spermatozoa, which were characterized by progressive motility and low-medium speed, as defined by high LIN and STR but low VCL, VSL and VAP values. Subpopulation 2 (SP2) was constituted by 14.72% of the total motile spermatozoa, defined by having the highest velocity and progressiveness, determined by high VCL, VSL, VAP, LIN and SRT values. Subpopulation 3 (SP3) included 23.78% of the total motile spermatozoa, characterized by hyperactivated-like movement (high speed but scarce progression), as defined by high VCL, ALH, and BCF, and low LIN and SRT. Finally, Subpopulation 4 (SP4) contained 37.22% of the total motile spermatozoa, including poorly motile and non-progressive spermatozoa, determined by low values for VCL, VSL, VAP, ALH, and BCF. Their distribution differed between experimental groups (Figure 2). For Group A, the distribution of the percentage of spermatozoa among the four different subpopulations with respect to the total count of spermatozoa was 15.97 ± 5.59, 9.88 ± 5.88, 17.37 ± 8.58, and 26.96 ± 8.23 for SP1, SP2, SP3, and SP4, respectively. Regarding Group B, the distribution of the percentage was 13.21 ± 9.26, 9.41 ± 11.06, 10.58 ± 9.46, and 15.95 ± 9.57 for SP1, SP2, SP3, and SP4, respectively. Concerning Group C, the distribution of the percentage was 11.76 ± 7.15, 7.25 ± 7.37, 13.94 ± 11.13, and 18.83 ± 11.24, for SP1, SP2, SP3, and SP4, respectively. Finally, the distribution of the percentage in Group D was 5.11 ± 4.54, 3.32 ± 4.64, 5.04 ± 6.59, and 9.19 ± 8.79 for SP1, SP2, SP3, and SP4, respectively. Statistically significant differences were observed between groups A, B, C, and the Group D in SP1 and SP2 (p < 0.05). Regarding SP3, control group significantly differed from groups A and C, and Group A significantly differed from Group B (p < 0.05). With respect to SP4, statistically significant differences were observed between Group D and groups A and C; similarly, Group A significantly differed from groups B and C (p < 0.05).

**Figure 2:**
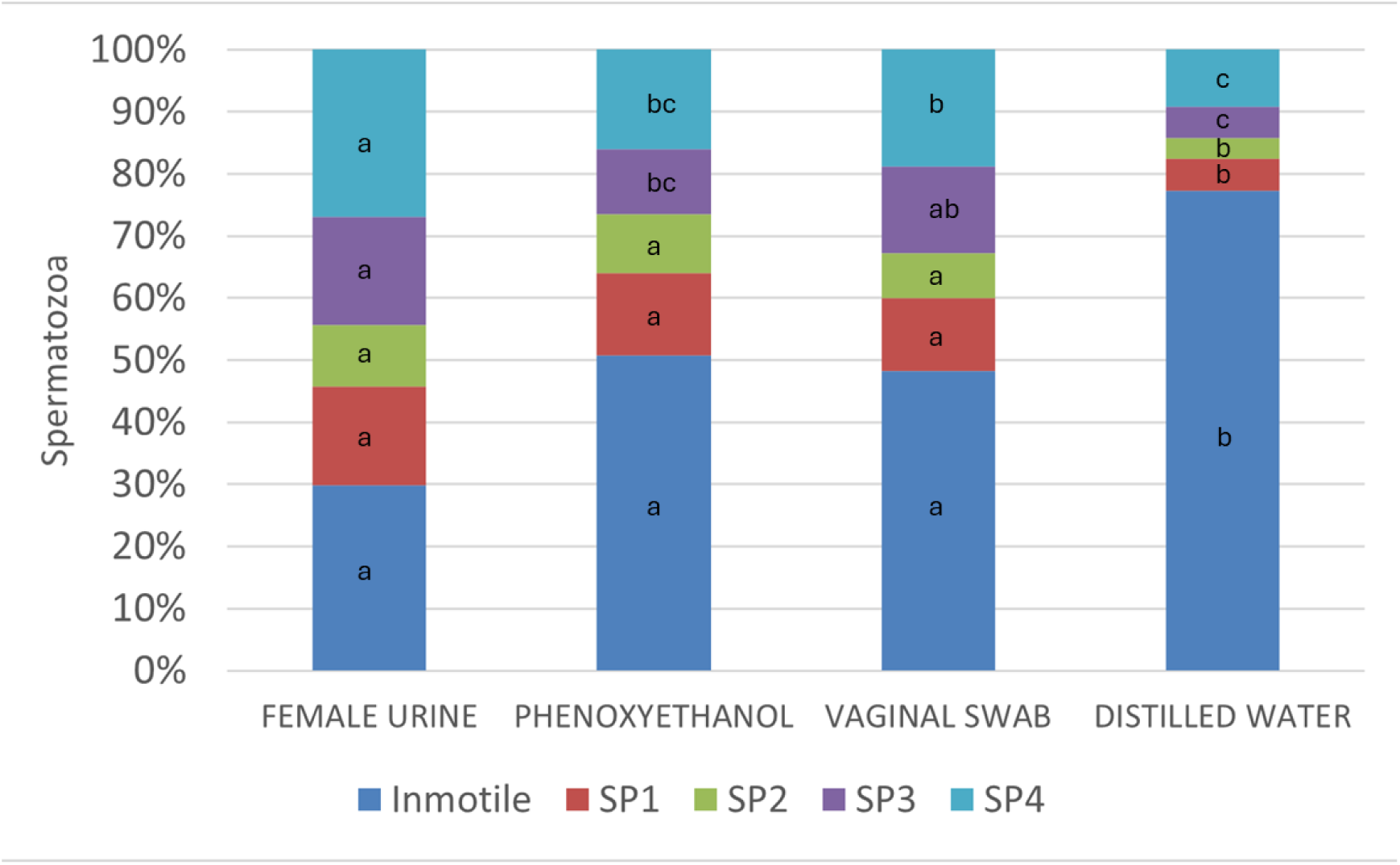
Distribution of the immotile and the four motile sperm subpopulations identified from sixty seven semen samples collected from male breeder rabbits after being exposed to doe urine, 2-phenoxyethanol, vaginal swab or distilled water (control) –potential sources of chemical communication-. Samples were preserved at 16°C and analysed using the CASA system ∼8 hours after collection. SP1: progressive sperm with low-medium speed; SP2: progressive sperm with high speed; SP3: spermatozoa with hyperactivated-like movement and low progressiveness; SP4: poorly motile non-progressive sperm. Different letters among equally coloured blocks indicate significant differences among columns (p < 0.05).

**Table 3:** Mean (± SD) kinematic parameters defining four motile sperm subpopulations identified from rabbit male breeder semen samples preserved at 16°C and analysed within 8 h after collection using the CASA system.

| Parameters | Subpopulations |  |  |  |
| --- | --- | --- | --- | --- |
|  | 1 | 2 | 3 | 4 |
| N° spz. (%) | 8937 (24.27) | 5420 (14.72) | 8757 (23.78) | 13705 (37.22) |
| VCL $\mu\text{m/s}$ | $45.51 \pm 20.75^{\text{a}}$ | $118.47 \pm 30.93^{\text{b}}$ | $130.24 \pm 32.26^{\text{c}}$ | $38.08 \pm 18.33^{\text{d}}$ |
| VSL $\mu\text{m/s}$ | $31.08 \pm 15.21^{\text{a}}$ | $80.03 \pm 23.22^{\text{b}}$ | $33.13 \pm 15.97^{\text{c}}$ | $6.60 \pm 5.51^{\text{d}}$ |
| VAP $\mu\text{m/s}$ | $35.36 \pm 15.71^{\text{a}}$ | $87.81 \pm 22.00^{\text{b}}$ | $64.58 \pm 16.63^{\text{c}}$ | $17.52 \pm 10.12^{\text{d}}$ |
| LIN % | $70.18 \pm 19.10^{\text{a}}$ | $69.93 \pm 18.88^{\text{a}}$ | $26.23 \pm 12.22^{\text{b}}$ | $17.52 \pm 12.05^{\text{c}}$ |
| STR % | $87.06 \pm 11.03^{\text{a}}$ | $90.93 \pm 8.79^{\text{b}}$ | $52.32 \pm 23.36^{\text{c}}$ | $37.81 \pm 23.04^{\text{d}}$ |
| WOB % | $79.80 \pm 15.54^{\text{a}}$ | $76.31 \pm 16.31^{\text{b}}$ | $50.40 \pm 10.13^{\text{c}}$ | $46.04 \pm 15.86^{\text{d}}$ |
| ALH $\mu\text{m}$ | $1.89 \pm 0.96^{\text{a}}$ | $3.39 \pm 1.81^{\text{b}}$ | $5.51 \pm 1.54^{\text{c}}$ | $2.17 \pm 0.97^{\text{d}}$ |
| BCF Hz | $4.05 \pm 4.14^{\text{a}}$ | $8.28 \pm 5.40^{\text{b}}$ | $9.10 \pm 4.68^{\text{c}}$ | $3.37 \pm 2.59^{\text{d}}$ |

Different superscript letters indicate significant differences between subpopulations ( $p < 0.001$ , except for VSL between subpopulations 1 and 3, $p < 0.002$ ). N° spz.: number of spermatozoa; VCL: curvilinear velocity; VSL: straight-line velocity; VAP: average path velocity; LIN: linearity; STR: straightness; WOB: wobble coefficient; ALH: mean amplitude of lateral head displacement; BCF: beat cross frequency.

### 3.3. Testosterone concentration

In addition to the sperm evaluation, we also characterized the levels of testosterone in male breeders, a primary male hormone that is involved in sexual receptivity and libido. ELISA assays were performed in five different blood serum samples per individual, taken at different time-points of the stimuli exposure, always before SC (Figure 1B). Testosterone concentration increased from the first time-point (prior to exposure) to the second time-point (1 h after the first exposure) in all experimental groups. Then, in groups C and D, it decreased for the rest of the time-points. However, in groups A and B, it decreased from the second time-point to the fourth time-point, and then increased again in the fifth time-point, especially in group A. Animals in group D (distilled water, control) showed the lowest testosterone concentration for the last two time-points. Overall, although not significant (p > 0.05), we found higher testosterone concentrations in males belonging to Groups A and C, compared to males exposed to groups B and D (Table 4, Figure 3).

**Figure 3:**
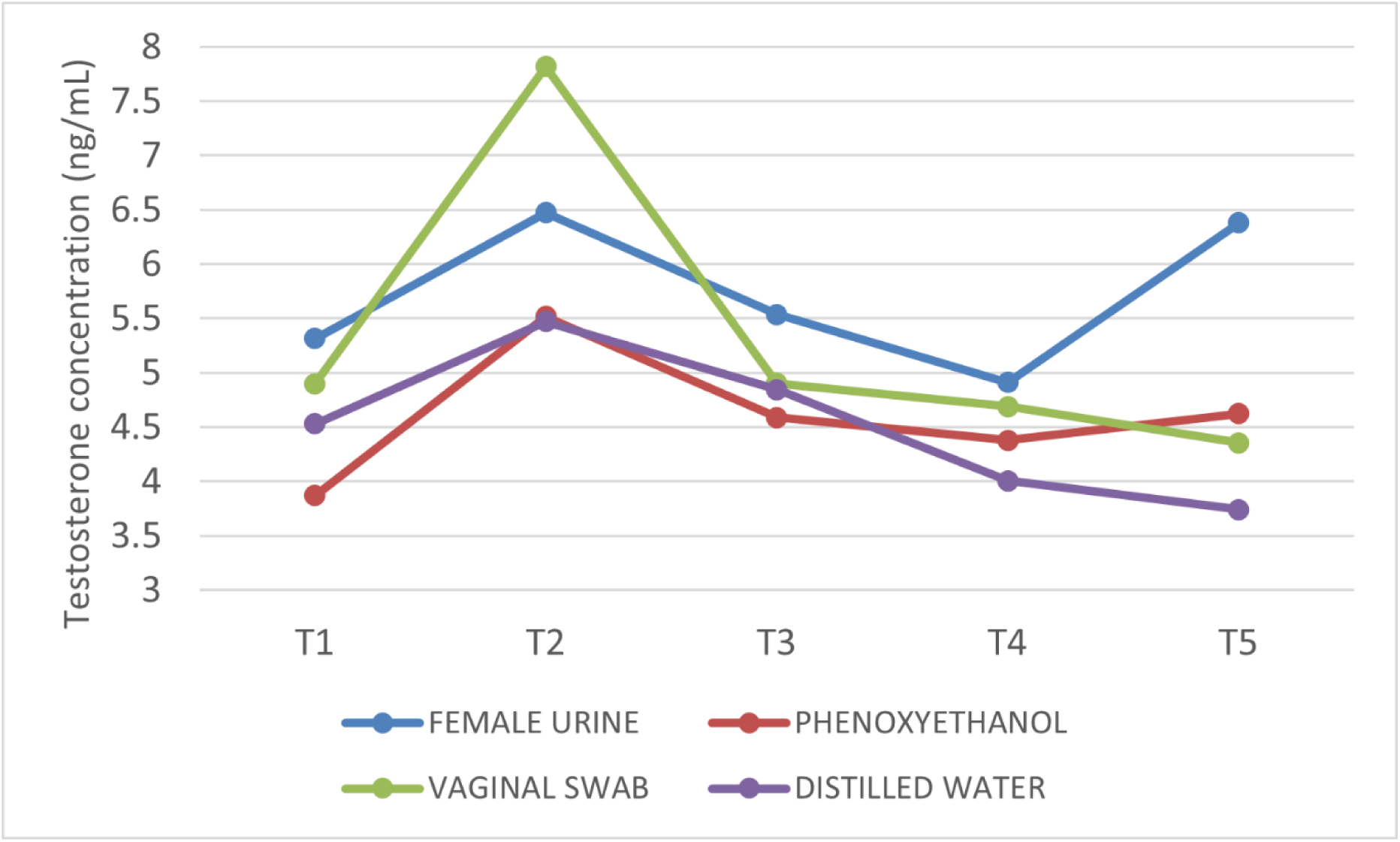
Graphical representation of the mean testosterone concentration (ng/mL) determined by ELISA analysis in male breeder rabbit serum samples, collected from 24 male rabbits before (T1) and during (T2, T3, T4, T5) exposure to female urine, 2-phenoxyethanol, vaginal swab and distilled water (control). T1: 150 min; T2: 90 min; T3: 60 min; T4: 30 min; T5: 2 min before semen collection.

**Table 4:** Mean (± SD) testosterone concentration (ng/mL) determined by ELISA analysis in male breeder rabbit serum samples collected from 24 male rabbits before (T1) and during (T2, T3, T4, T5) exposure to female urine, 2-phenoxyethanol, vaginal swab and distilled water (control). T1: 150 min; T2: 90 min; T3: 60 min; T4: 30 min; T5: 2 min before semen collection.

| Experimental group | Sample collection time |  |  |  |  |
| --- | --- | --- | --- | --- | --- |
|  | T1 | T2 | T3 | T4 | T5 |
| (A) Female urine | 5.31 ± 2.46 | 6.47 ± 4.07 | 5.54 ± 2.66 | 4.91 ± 3.31 | 6.38 ± 2.59 |
| (B) 2-phenoxyethanol | 3.87 ± 2.10 | 5.52 ± 2.14 | 4.59 ± 2.99 | 4.38 ± 2.25 | 4.62 ± 2.26 |
| (C) Vaginal swab | 4.90 ± 2.98 | 7.82 ± 6.81 | 4.90 ± 2.88 | 4.69 ± 3.01 | 4.36 ± 2.86 |
| (D) Distilled water | 4.53 ± 2.53 | 5.47 ± 3.02 | 4.36 ± 2.39 | 4.00 ± 1.49 | 3.74 ± 1.56 |

## 4. Discussion

We found higher percentage of motile, progressive, fast progressive and mid-progressive spermatozoa in males that were exposed to female biological fluids (doe urine and vaginal fluid) and 2-phenoxyethanol as a potential pheromone, compared to control individuals (exposure to distilled water). In contrast, semen volume and the percentage of immotile and non-progressive spermatozoa followed an opposite trend, being higher in the control group. We have also characterized four different sperm subpopulations as an additional source of comparison, and a higher percentage of fast and progressive spermatozoa was observed in all experimental groups compared to control group. Therefore, higher sperm motility was observed upon exposure to specific biological fluids potentially containing pheromones. Additionally, we found no differences in testosterone levels between groups, although males exposed to natural female biological fluids (either urine or vaginal fluid) showed a tendency towards higher testosterone levels.

### 4.1. Biostimulation using female biological fluids may increase semen quality in rabbits

It has been stated that the efficiency of AI relies on both male and female reproductive parameters. Regarding the male, success hinges on the effective generation of viable sperm, influenced by factors like the quantity of spermatozoa produced per ejaculation and the quality attributes associated with its fertilization potential [29]. In this way, the percentage of motility has been pointed out as a determinant for fertility, prolificacy, and productivity [5], as the higher the number of motile spermatozoa in the AI dose, the higher the female productivity. Similarly, [29] observed a positive relationship between sperm motility parameters and productivity, with an increase of 1.1 live kits/litter when the percentage of motile sperms exceeded 83%. According to our results, groups A, B, and C, exposed to female urine, 2-phenoxyethanol, and vaginal fluid, respectively, showed higher motility levels than group D, exposed to distilled water (control). Therefore, we speculate that higher male productivity could be achieved by exposing the males to biological compounds likely containing pheromones, before SC. Nevertheless, it should be considered that fertility encompasses several factors alongside semen traits (such as female management and health status, and proper AI technique), which must be considered when dealing with reproductive efficiency in farms.

To perform a complete semen analysis, we determined the subpopulations present in the seminal doses. This kind of analysis is based on individual sperm kinetics, and it has been reported in animal species such as cats, dogs, stallions, sheep, boars, bull, and rabbits, among others [30–36]. Indeed, the presence of such structure in mammals spanning diverse phylogenetic origin may imply a potential correlation between alterations in the specific subpopulation arrangement within an ejaculate and its capacity for fertilization [37]. Accordingly, we considered it important to elucidate the possible changes on motile sperm subpopulations that could be induced following exposure to secretions that may contain pheromones. In our experiment, statistically significant differences were obtained regarding Subpopulation 2 (progressive sperm with high speed) between groups A, B, C and group D. It has been previously observed that fast and progressive sperm correlates with better fertilization rates [38]. Therefore, as it has been mentioned above, higher productivity could be expected when using pheromone-like stimulation prior to SC. To our knowledge, this is the first study to perform a detailed semen analysis of samples collected from male breeder rabbits exposed to biological compounds potentially containing pheromones, and therefore no comparisons with other data sources can be discussed. In this way, an increase in semen quality would have potential benefits also in female reproduction. Further studies should evaluate reproductive parameters, such as fertility and prolificacy rates (following the protocol described in [14]) of female rabbits when they are inseminated with semen of males exposed to female urine, vaginal secretion or 2- phenoxyethanol. Little increases in fertility and/or prolificacy would be highly beneficial for the farm management and the competitiveness of the sector.

As for reaction time, no differences between groups were observed, in contrast to other studies in which male breeders were exposed to does prior to SC [26]. One possible explanation could be the need of visual and/or physical stimuli to increase libido (e.g. actual presence of the doe, not only exposure to biological compounds). However, it should be considered that libido also depends on other factors such as age. Some of the animals that we used were over three years old, and the literature points to a decrease in reproductive efficiency after 2 years old [10]. Other considerations such as reproductive rhythm, housing, feeding, genotype, and photoperiod may also influence male reproductive parameters [26,39,40].

### 4.2. The potential role of 2-phenoxyethanol in rabbit reproduction

We employed 2-phenoxyethanol, a potential chemical stimulus to enhance male reproductive parameters. This compound has been previously identified in chin gland secretions of dominant, but not subordinate, adult male rabbits [23,41], where it functions as a fixative to preserve dominant scents in the environment. Other compounds known as major urinary proteins, have been found to serve as both fixatives and pheromones in mice, depending on social context [42,43]. In the literature, examples of 2-phenoxyethanol being utilized as a pheromone are scarce and confined mainly to insects. For instance, 2-phenoxyethanol present in ball-point pen ink acts as an analogue of trail pheromone in termites [44], and the same molecule has been identified as a pheromone in burying beetle, implicated in parental behaviour triggering offspring begging [24]. In our study, given the absence of characterized pheromones associated with sexual behaviour in rabbits, we employed 2-phenoxyethanol to explore its potential as chemical stimulus that may enhance male reproductive parameters. By exposing male rabbits to 2-phenoxyethanol, we observed an increase in semen quality but not quantity, suggesting a potential role of this compound in improving reproductive efficiency. The source of 2- phenoxyethanol, whether it is released during male-male interactions or emitted as a stimulant by female rabbits, remains uncertain. Future experiments are necessary to clarify the origin and function of this chemical compound in the context of reproductive efficiency in rabbits, including examining whether it is emitted by males during encounters or originates from the female chin gland. Furthermore, prior to implementing it as a biostimulation method, further studies should account for an increase in the sample size (to assess any potential effects on semen volume) or explore its effects on female reproduction. Lastly, we established an experimental framework that facilitates future studies on pheromone identification for biostimulation purposes in rabbit husbandry.

### 4.3. Importance of testosterone in male reproductive efficiency

In male mammals, testosterone is released in a pulsatile manner alongside low “basal” levels. Depending on the context, two distinct types of testosterone release occur: spontaneous and reflexive. Spontaneous release dominates male circulating testosterone, occurring multiple times daily in response to internal homeostatic conditions, crucially shaping various aspects of male reproductive physiology and behaviour. Conversely, reflexive testosterone release is less frequent, triggered by mating and aggressive interactions often involving pheromones [45–47]. In fact, pheromone sensing stimulates the release of testosterone, and this hormone impacts neural and behavioural responses in a sex-specific manner [48,49]. At functional levels, the swift elevation in testosterone levels during these social behaviours suggests a role for non-genomic mechanisms in converting testosterone to oestradiol (E2) via aromatase [50]. While the precise cues and physiological processes governing reflexive testosterone release remain unclear, sexual cues have been observed to boost testosterone levels [51,52], and semen quality and volume [52,53]. For instance, exposure to female vaginal secretions triggers increased plasma LH and testosterone in male hamsters [54], and testosterone quickly enhances ejaculate volume and sperm density in male goldfish [50]. However, it is worth noting that different species exhibit varying responses; for instance, chemosensory cues have no effect on testosterone levels in male terrestrial salamanders [54]. Moreover, the role of plasticity in testosterone function adds another layer of complexity, allowing individuals to adapt their behavioural responses to environmental changes [47].

In our study, we observed a rise in testosterone levels at the second time-point compared to the first, followed by a decline after the third time-point. This suggests a potential increase in testosterone levels via the reflexive mechanism upon initial exposure to the stimulus. We hypothesize that the subsequent decrease in testosterone levels could be associated with social experience and functional plasticity. Additionally, although our findings lacked statistical significance (p > 0.05), we speculate that female fluids (doe urine and vaginal swab) may influence male testosterone concentration, potentially boosting male receptivity, libido, and semen quality and volume. Further experiments increasing the number of animals or analysing testosterone just after exposure to the fluids will help confirm our hypothesis.

### 4.4 Doe urine and vaginal fluid as potential biostimulators

Urine and vaginal fluid have been widely recognized as significant sources of pheromones across numerous species, attributed to their capacity to modulate male sexual behaviour and reproductive functions [55–57]. While previous investigations have indicated that exposure of male rabbits to females elicits heightened sexual drive and improved semen characteristics [26], a comprehensive assessment of the specific female cues driving these alterations has not been undertaken. In our study, we exposed male rabbits to doe urine or vaginal secretions as potential sources of pheromones, and we demonstrated a pronounced enhancement in semen quality and a trend towards elevated testosterone levels in rabbit males upon exposure to both female biological fluids. Although our findings are preliminary and require further validation, they collectively suggest the potential involvement of these secretions as pheromone releasers. Therefore, further studies in this field should prioritize elucidating the biochemical properties of these compounds with the objective of identifying putative pheromone-like substances amenable to experimental validation. Additionally, it is important not to overlook other potential biological sources of pheromones, such as exocrine glands. For instance, the chin gland has been identified as particularly relevant for rabbit social behaviour [19]. In light of this, we suggest that extracts from female glands could be exposed to males to investigate their possible contribution to reproduction. Moreover, employing analytical techniques such as HPLC or mass spectrometry could facilitate the identification of the molecular identity of potential pheromone candidates. Characterizing the molecular nature of sex pheromones in rabbit farming holds promise for the industry, offering the prospect of integrating pheromones, as natural cues, into routine husbandry practices for potential enhancement of reproductive outcomes. Such integration would ultimately contribute to fostering a more natural, organic, and respectful image of the sector.

## 5. Conclusions

Our findings suggest that exposure to female urine, vaginal fluid, and 2-phenoxyethanol as a potential reproductive pheromone, enhance rabbit sperm quality, thereby underscoring the significance of biostimulation methods based on chemical communication in rabbit farming. Moreover, while blood testosterone levels did not significantly differ among groups, males exposed to natural female biological fluids (either urine or vaginal fluid) exhibited higher testosterone levels. Future experiments should not overlook testosterone as a marker when assessing pheromone-mediated reproductive efficiency and should investigate the molecular composition of body secretions potentially containing pheromones, as well as assess other biological fluids or exocrine glands. This research will be essential to facilitate the integration of pheromones into rabbit husbandry practices, thereby improving both animal production and welfare and transitioning toward a more sustainable and organic system.

## CRediT authorship contribution statement

**Paula R. Villamayor**: Conceptualization, Methodology, Validation, Investigation, Writing – original draft preparation, Supervision. **Uxía Yáñez**: Methodology, Validation, Investigation, Formal analysis, Data curation, Writing – original draft preparation. **Julián Gullón**: Validation, Resources, Investigation, Visualization, Writing – review and editing. **Pablo Sánchez-Quinteiro**: Conceptualization, Resources, Writing – review and editing, Visualization, Project administration, Funding acquisition. **Ana I. Peña:** Investigation, Formal Analysis, Data Curation, Visualization. **Juan José Becerra:** Investigation, Validation, Visualization, Writing – review and editing**. Pedro G. Herradón:** Investigation, Validation, Visualization, Writing – review and editing. **Paulino Martínez:** Conceptualization, Resources, Writing – review and editing, Visualization, Supervision. **Luis A. Quintela**: Conceptualization, Formal analysis, Resources, Writing – review and editing, Visualization, Supervision, Project administration, Funding acquisition. All authors have read and agreed to the published version of the manuscript.

## Acknowledgements

This publication is part of the I+D+i proyect grant PID2021-127814OB-I00 funded by MCIN/AEI/10.13039/501100011033 and by “ERDF A way of making Europe”. PRV and UY were supported by research fellowships funded by Xunta de Galicia (refs. ED481A- 2020/491430, and 2020/122, respectively). The authors thank COGAL SA. (Pontevedra, Spain) for providing the facilities and animals employed in this study, as well as technical support. We also gratefully thank Adrián Casanova, Marina Pampín, Ahlam Zahourti, Isabel Cavalcanti, Jennifer Ríos, and Patricia Regal for their technical support.

